# MYC hyperactivates WNT signaling in APC/CTNNB1-mutated colorectal cancer cells through miR-92a-dependent repression of DKK3

**DOI:** 10.1101/2021.07.26.453875

**Authors:** Priyanka Sehgal, Claudia Lanauze, Xin Wang, Katharina E. Hayer, Manuel Torres-Diz, Yogev Sela, Ben Z. Stanger, Christopher J. Lengner, Andrei Thomas-Tikhonenko

## Abstract

Activation of Wnt signaling is among the earliest events of colon cancer development. It is achieved either via activating mutations in the CTNNB1 gene encoding β-catenin, the key transcription factor in the Wnt pathway, or most commonly by inactivating mutations in APC, a major β-catenin binding partner and negative regulator. However, our analysis of recent Pan Cancer Atlas data revealed that CTNNB1 mutations significantly co-occur with those affecting Wnt receptor complex components (e.g., Frizzled and LRP6), underscoring the importance of additional regulatory events even in the presence of common APC/CTNNB1 mutations. In our effort to identify non-mutational hyperactivating events, we determined that KRAS-transformed murine colonocytes overexpressing direct β-catenin target MYC show significant upregulation of the Wnt signaling pathway and reduced expression of Dickkopf 3 (DKK3), a reported ligand for Wnt co-receptors. We demonstrate that Myc suppresses Dkk3 transcription through one of mir-17-92 cluster microRNAs, miR-92a. We further examined the role of DKK3 by overexpression and knockdown and discovered that DKK3 suppresses Wnt signaling in APC-null murine colonic organoids and human colon cancer cells despite the presence of downstream activating mutations in the Wnt pathway. Conversely, MYC overexpression in the same cell lines resulted in hyperactive Wnt signaling, acquisition of epithelial-to-mesenchymal transition markers, and enhanced migration and invasion and metastasis in syngeneic orthotopic mouse colon cancer model. Our results suggest that the MYC->miR-92a-|DKK3 axis hyperactivates Wnt signaling, forming a feedforward oncogenic loop.

**SIGNIFICANCE:** Common APC and CTNNB1 mutations activate Wnt signaling in colorectal cancers. Here we demonstrate that further potentiation of this pathway involves microRNA-dependent repression of the DKK3 gene by the Myc oncoprotein.

## INTRODUCTION

Colorectal cancer (CRC) is the second most common cancer diagnosed and third leading cause of cancer-related deaths in the United States. Despite serving as the initial testing ground for cancer genetics and more recently - genomics, CRC remains a deadly disease. There are projected to be 147,950 individuals newly diagnosed with CRC with an estimated 53,200 CRC deaths in 2020 (1). One reason for the lack of major breakthroughs is that focusing on individual signaling pathways is not enough to understand pathogenesis and progression of this disease. The MYC oncogene is a case in point. It is involved in a dizzying number of functional interactions, few of which have been fully understood or sufficiently validated.

In CRC, MYC is usually overexpressed due to activating mutations in the WNT pathway (2,3). Binding of WNT ligand to its receptor Frizzled (FZD) and co-receptor Low density lipoprotein receptor related protein 5/6 (LRP5/6) initiates a sequence of signaling events, which reprograms protein-protein interactions and ultimately prevent Adenomatous Polyposis Coli (APC) tumor suppressor-mediated degradation of β-catenin (4,5). Stabilized β-catenin translocates into the nucleus (6), where it forms a complex with the TCF4 transcription factor (7,8) and drives expression of MYC, along with numerous other target genes involved in stem cell self-renewal and proliferation (3). Not surprisingly, inactivating mutations in APC and activating mutations in CTNNB1 (which encodes β-catenin) are mutually exclusive as they work towards the same goal of rendering Wnt signaling pathway constitutively active (9,10). The question remains whether APC- or CTNNB1-mutant CRCs accumulate complementary genetic or regulatory events that are needed to hyperactivate Wnt signaling.

In this study we set out to identify such hyperactivating events by establishing epistatic relationships between various recurrent mutations in the Wnt pathway in CRC. We discovered that mutations of many additional components of the Wnt signaling co-occur with CTNNB1 mutations, consistent with our “Wnt hyperactivation” hypothesis. These mutations are exemplified by the 4 members of the DKK superfamily, which encode secreted glycoproteins capable of modulating the activity of LRP5/6 and Kremen co-receptors in cells without genetic perturbation of the Wnt pathway (11-13). We then focused on the least frequently mutated member in the Wnt signaling pathway: DKK3, whose role in Wnt signaling (13,14) and CRC is not well defined. We discovered that although it is seldom mutated, it is negatively regulated by Myc by a microRNA-dependent mechanism and that WNT, Myc, miR-92a and DKK3 form a previously unrecognized positive feed forward loop wherein Myc and Wnt activate each other.

## MATERIALS AND METHODS

### Cell culture

The human CRC cell lines HCT116, SK-CO-1, mouse colonocytes Ras and RasMyc, Mouse L cells with or without Wnt3a were cultured in DMEM (Invitrogen) supplemented with 10 % fetal bovine serum (FBS; Invitrogen), 2mM L-glutamine, penicillin/streptomycin (p/s) at 37°C and 5% CO2. After thawing, cells were authenticated by short tandem repeat analysis, tested for mycoplasma using the EZ-PCR Mycoplasma PCR Detection Kit (Biological Industries), and used for up to 12 passages. Mouse APC null, p53 null, Kras mutant intestinal organoids were maintained in DMEM-F12 supplemented with 10 % fetal bovine serum, 2mM L-glutamine, penicillin/streptomycin (p/s), N2 supplement, B27 supplement and N-acetyl-cysteine.

### Organoid generation

The colonic crypts from a KrasG12D-LSL mouse (The Jackson Laboratory, stock no: 008179) were extracted and used to establish colonoid cultures. The KrasG12D mutation was then activated by transient transfection of Salk-Cre with pPGK-Puro (Addgene#11349) plasmids, followed by puromycin selection for 3 days. Next, Apc, p53, Smad4 mutations were introduced by CRISPR-Cas9 editing. Specifically, sgRNAs targeting Apc, p53 and Smad4 were cloned into PX330 plasmid (Addgene#42230) and transiently transfected into the KrasG12D tumoroids. One week after the transient transfection, the tumoroids with Apc, p53, and Smad4 mutations were selected by removing R-spondin, adding Nutlin-3 and removing Noggin from the culture media, respectively. 10 subclones were picked from the engineered bulk tumoroids, conditional PCR and Sanger sequencing were used to verify the mutations in each subclone. A subclone with the recombined LSL-KrasG12D allele, and verified Apc, p53, and Smad4 mutations was used for downstream experiments.

### RNA extraction and quantitative real-time PCR

Total RNA, including miRNA, was isolated from tissues or cell lines using TRIzol reagent (Invitrogen) according to manufacturer’s instructions. For DKK3 and Myc mRNA detection and miRNA expression analysis, reverse transcription was performed using the ABI cDNA reverse transcription kit (Applied Biosystems) with miR-92a-specific primers (Applied Biosystems). Quantitative PCR was performed using FAST SYBR green mix (Roche) on the ABI 7500 real-time PCR System (Applied Biosystems). U6 snRNA or HPRT was used as internal control. The primer sequences are available upon request. The relative expression levels were calculated by the equation 2-ΔΔCT.

### Western blot analysis

Total cell lysates were prepared from cultured cells or organoids using RIPA buffer with protease and phosphatase inhibitors (Pierce Halt Inhibitor Cocktail, Thermo Scientific). After protein transference to PVDF (Immobilin-p, Millipore), the antibodies for p-LRP6, LRP6, AXIN2, CCND1, Myc, GSK3β, p-GSK3β (Ser9), DKK3, Phospho β-catenin, β-catenin (Cell Signaling Technology) were used. Subsequently, recommended dilutions of HRP conjugated secondary antibodies (GE healthcare) were applied. Enhanced chemiluminescence (ECL; Millipore) was used to detect bands and captured by Chemiluminescence imager (GE healthcare). Each sample was normalized to GAPDH or β-actin (Cell signaling).

### Plasmids and transfection

For infection of CRC cell lines with pMX-IRES constructs, retroviral particles were generated by transfection of HEK 293GP cells with Lipofectamine 3000 (Invitrogen). After transfection, the cell supernatants were collected and used to infect CRC cells, and the stably transfected cells were selected using puromycin and confirmed by quantitative RT-PCR. Selection for infected cells was done with 12.5 ug/ml Blasticidin (Gemini Bio Products) for over a week. SMARTpool siRNA for DKK3, Antagomir-92a, miR-92a mimic and corresponding control oligonucleotides (Dharmacon) were transfected into CRC cells by using Lipofectamine 3000 according to manufacturer’s protocols. For stable infection of organoids, PMX-IRES GFP constructs were used. The retroviral particles were prepared as mentioned above and concentrated using reterconcentrator (Takara Biosciences). Organoid fragments were prepared following the protocol by (15) and combined with retroviral solution along with polybrene in a 24-well plate, sealed, and spinoculated at 1800 rpm at 37°C for 1 hr 45 min. Following spinoculation, plate was incubated for 6 hrs at 37°C. This was followed by seeding of infected organoids and 5 days post infection, GFP positive cells were sorted and resuspended in matrigel.

### Luciferase reporter assay

Cells were transfected with either TOP flash (containing TCF binding sites; Addgene#12456) or FOP flash (mutated TCF binding sites; Addgene#12457) luciferase expression vectors and with Renilla luciferase vector (control) using Lipofectamine 3000. 24 hours post-transfection, cells were incubated with Wnt3a CM. 20 hrs post transfection, luciferase activity in total cell lysates was measuring using Dual-Luciferase Reporter Assay Kit (Promega) and normalized for transfection efficiency via dividing by the Renilla luciferase activity. The TOP/FOP ratio was used as a quantitative measure of β-catenin-mediated transcriptional activity as described by us recently (16). To validate whether DKK3 is a direct target of miR-92a, wild type or mutant 3′-UTR of DKK3 was cloned into the psicheck-2 vector (Promega). HCT116 cells were transfected with and wild-type or mutant 3′-UTR-luc by using Lipofectamine 3000. After transfection for 48h, cells were harvested and assayed with Dual-Luciferase Reporter Assay System (Promega) according to the manufacturer’s protocols. The sequences of primers used for generating wild type or mutant 3’-UTR are available upon request.

### Biotin-labeled microRNA pull-down assays

Biotinylated miR-92a (Dharmacon) pull-down assay with target mRNAs was performed as described earlier (17). Briefly, 1 × 106 HCT116 cells were seeded in 10 cm plate 24 hrs later, control miR or 3’ biotin-labeled miR-92a was transfected at a final concentration of 50 nM. After 24 hours, whole cell lysates were harvested. Streptavidin-Dyna beads (Dyna beads M-280 Streptavidin, Invitrogen, 50 μl each sample) were coated with 10 μl per sample yeast tRNA (stock 10 mg/ml Ambion, Austin, TX) and incubated with rotation at 4°C for 2 hrs. Then beads were washed with 500 μl lysis buffer and resuspended in 50 μl lysis buffer. Sample lysates were mixed with pre-coated beads (50 μl per sample) and incubated overnight at 4°C on a rotator. Beads were then pelletted down the next day to remove unbound materials at 4°C for 1 minute, 5K rpm and washed five times with 500 μl ice cold lysis buffer. To isolate the RNA, 500 ul of TRIzol (Invitrogen) was added to both input and pulldown samples. Tubes were mixed well and kept in −20°C for 2 hrs. RNA was then precipitated using standard chloroform-isopropanol method and then subjected to q-PCR.

### Migration and invasion assays

The Corning BioCoat Cell Culture Inserts and Matrigel inserts (Corning, USA) were used for migration and invasion assays, respectively. Briefly, the inserts were rehydrated with plain DMEM for 2 h before use. 5-7×10^4^ cells were trypsinised and resuspended in serum-free medium and then seeded onto 24-well transwell chambers with 8-μm pore membrane in 500 μL serum-free medium with condition medium from Wnt3A-producing L cells or control L cells. The lower chamber contained medium supplemented with 10% FBS. After incubation for 22 h, the non-migrated/invaded cells on the upper side of membrane were removed with a cotton swab and the migrated/invaded cells stained with crystal violet and photomicrographed.

### Orthotopic mouse model

Tumoroids harboring engineered oncogenic mutations in *Apc, p53, Kras* and *Smad4* (4×10^6^ cells) with or without Myc overexpression were implanted orthotopically into the cecal wall of syngeneic Bl/6 mice and allowed to engraft for 6 weeks. At that time, primary tumors and livers were resected and analyzed histologically to evaluate for metastatic lesions in the liver.

### Bioinformatics analysis

Publicly available data for “Colorectal Adenocarcinoma (TCGA, PanCancer Atlas)” was downloaded from cBioPortal (coadread_tcga_pan_can_atlas_2018.tar.gz). From these data we examined the mutual exclusivity of mutations and the normalized RNA-seq expression. We correlated Myc and DKK3 expression using R’s lm method to determine R2 and significance. Microarray (Array express database under accession number E-MEXP-757) intensities were normalized and analyzed using the Bioconductor package limma. The volcano plots only display the probes with the highest absolute fold change. Results were processed and visualized using the R packages dplyr and ggplot2.

### Statistical analysis

All statistical analyses were carried out using Graph Pad Prizm (version 7) by unpaired student’s t-test for two group comparisons or one-way ANOVA correcting for multiple comparisons, with similar variance between groups being compared. Error bars represent s.e.m. ± SD, and statistical significance was defined as *P* < 0.05.

## RESULTS

### Multiple Wnt pathway mutations co-occur in APC/CTNNB1-mutant CRC

In 2012, The Cancer Genome Atlas project (TCGA) reported genome-scale analysis of 276 samples, including exome sequence, DNA copy number, promoter methylation and messenger RNA and microRNA expression (18). In addition to APC and CTNNB1, that paper identified several key members of the Wnt pathway being mutated, including Frizzled (FZD10), LRP co-receptors, DKK 1-4, Axin2, FBXW7 and TCF7L2. In 2017 this dataset was supplemented with additional specimens resulting in the larger 594-sample CRC subset of TCGA PanCancer Atlas (19). We excluded from our analysis APC mutations given their very high prevalence (75%). The Oncoprint implemented on cBio portal provided updated frequency numbers (Figure 1A), and the mutual exclusivity analysis surprisingly revealed that all statistically significant relationship were in fact co-occurrences (Figure 1B). For example, CTNNB1 mutations co-occurred with those in LRP6, suggesting that even in β-catenin-mutant CRCs receptor-mediated events still play an important role. In turn, LRP6 mutations co-occurred with those in three members of the DKK family (1,2, and 4), suggesting the importance of deregulating both receptors and their cognate ligands. Of note, DKK3 did not appear in this list, casting uncertainty on its role in Wnt signaling and CRC pathogenesis in general. However, by analyzing its expression across various solid cancer types profiled in the Broad Cell Line Encyclopedia, we observed that CRC has the lowest DKK3 mRNA expression levels (Supplemental Figure S1A), suggesting that this least frequently mutated DKK family member is subject to additional mechanisms of dysregulation, such as gene repression.

**Figure 1.**
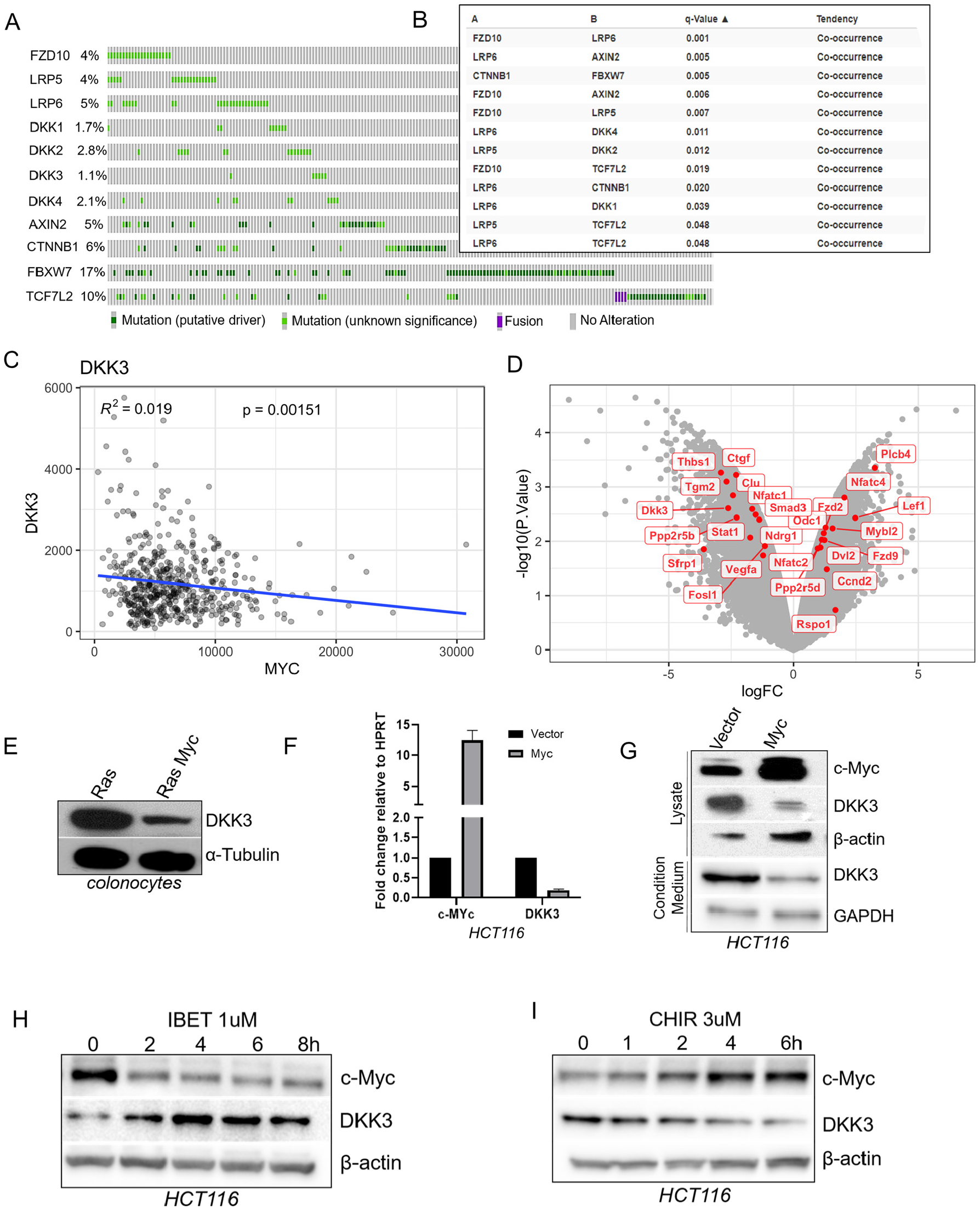
Myc regulates DKK3 expression in colon cancer cells. (A) Oncoprint analysis showing frequency of mutation in Wnt pathway members. (B) Mutual exclusivity analysis showing significant co-occurrence of mutations in Wnt pathway members. (C) Inverse correlation between Myc and DKK3 expression in human CRC. r=-0.14. (D) Volcano plot showing expression of TGFβ and Wnt pathway components in KRAS and RAS+MYC cells. (E) Immunoblotting on total lysates from the same cells performed using an anti-DKK3 antibody. α-tubulin was used as a loading control. (F) qRT-PCR analysis performed on parental and Myc-overexpressing HCT116 cells. Bar graph represents log2-transformed ratios of MYC and DKK3 expression normalized to HPRT. (G) Immunoblotting showing protein levels of Myc and DKK3 in the same cells. β-actin was used as a loading control. Also shown are levels of DKK3 in the conditioned medium. (H) Immunoblotting showing protein levels of Myc and DKK3 in HCT116 cells treated with IBET-151 (1 µM) for 0-8hrs. (I) Immunoblotting showing protein levels of Myc and DKK3 in HCT116 cells treated with Chiron-99021 (3 µM) for 0-6 hrs.

### Myc regulates DKK3 expression in colon cancer cells

To identify possible mechanisms of repression, we analyzed transcription factors-encoding mRNAs which anti-correlated with DKK3 mRNA in human samples from the TCGA CRC dataset. Interestingly, one of the transcription factors that showed significant anticorrelation with DKK3 was MYC (Figure 1C). In order to establish the causality, we utilized the previously generated MYC-dependent CRC model system from our lab (20). Specifically, we compared the transcriptomic profiles of MYC-overexpressing and parental KRAS-transformed mouse colonocytes. Volcano plot analysis of the Myc-upregulated genes yielded some of key Wnt pathway components (Lef1, Dvl2, Fzd2/9, etc; Figure 1D) indicating that overexpression of MYC alters Wnt signaling pathway. It also demonstrated lower expression levels of DKK3 along with several TGFβ-responsive genes reported by us previously (21), suggesting that MYC negatively regulates DKK3. Immunoblotting experiments confirmed that Myc-overexpressing colonocytes have reduced expression of DKK3 at the protein level (Figure 1E). To extend our findings to human colon cancer cells, we used the HCT116 derivative stably overexpressing Myc (22). We found that Myc-overexpressing cells have decreased expression of DKK3 mRNA (Figure 1F) and DKK3 protein levels were decreased in lysates as well as in the conditioned medium (Figure 1G), indicative of negative regulation of DKK3 by Myc.

To confirm the effect of Myc on DKK3 independently of retrovirus-mediated overexpression, we used chemical inhibitors (I-BET 151 and CHIR99021) to repress or stabilize Myc expression in HCT116 cells, respectively. First, using the chemical inhibitor of the Brd4 transcription factor, I-BET 151 (23), we were able to inhibit Myc expression and observed a concomitant increase in DKK3 levels at corresponding time intervals (Figure 1H). Using a reverse strategy, we stabilized Myc levels in HCT116 cells using CHIR99021 treatment, as described by us previously (24). CHIR99021 inhibits the phosphorylation of Myc at Threonine-58 by GSK-3-β, which leads to phospho-Myc being targeted for ubiquitination and subsequent degradation (25). Myc levels thus stabilized resulted in decreased DKK3 expression (Figure 1I), confirming an inverse relationship between Myc and DKK3.

### Myc regulates DKK3 expression via miR-92a

To further understand the regulation of DKK3 expression by MYC, we mined the publicly available dataset of Myc chromatin immunoprecipitation (ChIP) from the colon cancer cell line HCT116 [GEO dataset GSM2576763 from (26)]. We observed that MYC does not bind to the promoter of DKK3 as evidenced by the absence of peaks within 3Kb of the transcription start site. This is in contrast to other genes reported to be directly repressed by Myc, such as NDRG1 (27) (Figure 2A). This negative result was indicative of an indirect mechanism of regulation, for example by a Myc-regulated microRNA (28).

**Figure 2.**
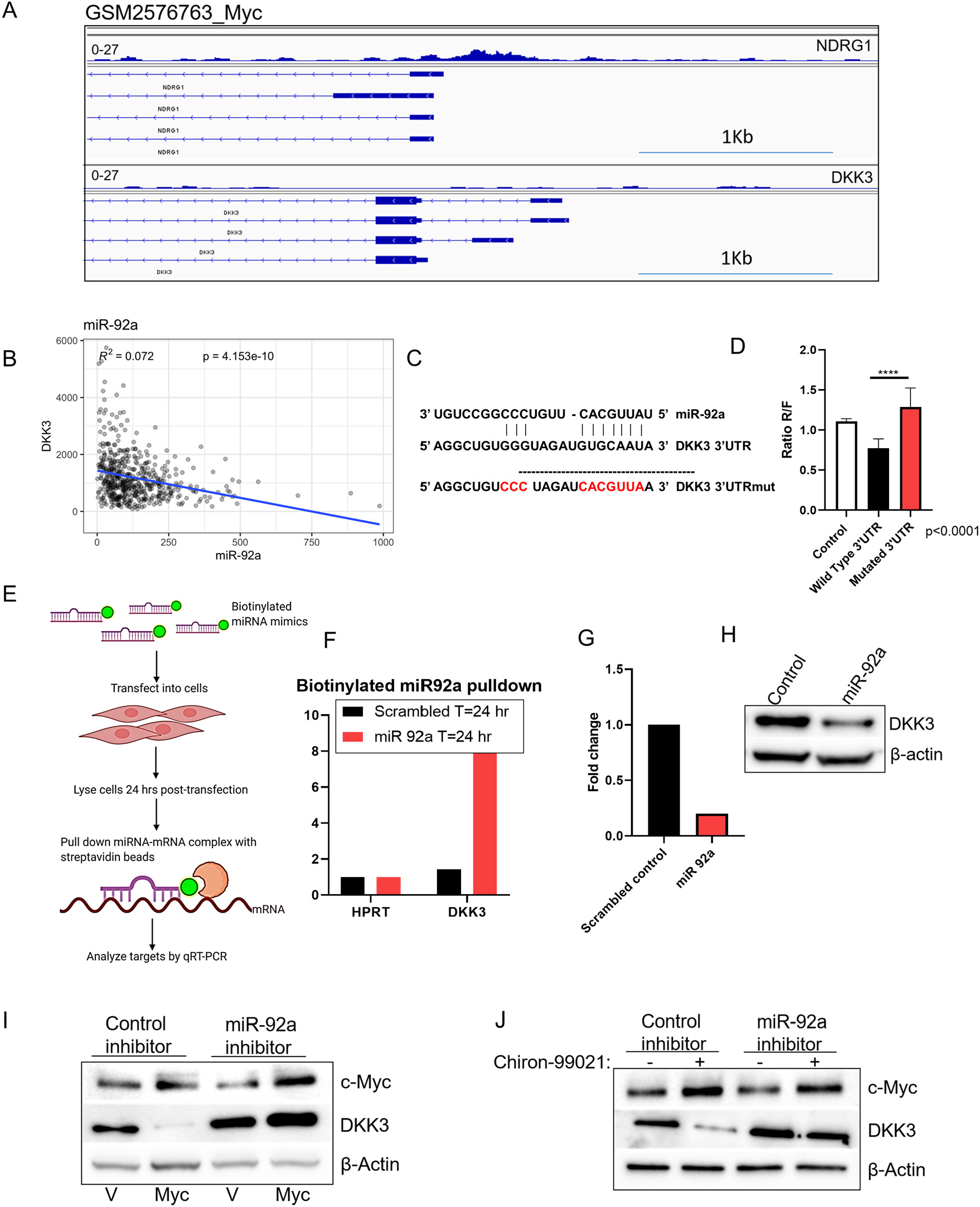
Myc regulates DKK3 expression via miR-92a in CRC cells. (A) CHIP Seq analysis of NDRG1 and DKK3 promoters for Myc binding. (B) Inverse correlation between miR-92a and DKK3 expression in human CRC. r=-0.26. (C) Nucleotide sequence of wild type and mutant 3’ UTR of DKK3. (D) Luciferase activity of DKK3 wild type and mutant 3’-UTR -base reporters in HCT116 cells. (E) Schematic for biotinylated miR-92a pull down with streptavidin beads. (F) Enrichment for DKK3 mRNA in streptavidin pull-downs from cells transfected with biotinylated miR-92a mimic. HPRT transcript was used as a control. (G) Quantitation of DKK3 mRNA expression normalized to HPRT (log2-transformed ratio) by qRT-PCR analysis in control and miR-92a transfected cells. (H) Immunoblotting showing level of DKK3 in the same cells. GAPDH was used as loading control. (I) Quantitation of miR-92 effects on DKK3 levels by immunoblotting. Vector-transduced and Myc-overexpressing HCT116 cells were treated with control or miR-92a inhibitor. Total cell lysates were probed for Myc and DKK3 as indicated. β-Actin was used as loading control. (J) Quantitation of miR-92 effects on DKK3 levels by immunoblotting. Parental HCT116 cells were treated with control or miR-92a inhibitor in the presence or absence of Chiron-99021 (3 µM for 6 hrs). Total cell lysates were probed for Myc and DKK3 as indicated. β-Actin was used as loading control.

Previous data by several labs, including ours, has shown that Myc upregulates miR-17-92 cluster of microRNA (20,29,30). Per TargetScan (31), one of the predicted targets for miR92a is DKK3 (Supplementary Figure S1B). Accordingly, we observed highly significant anti-correlation between miR-92a and DKK3 mRNA in human samples from the TCGA CRC dataset (Figure 2B). To further confirm that DKK3 is a target of miR-92a, we generated WT-DKK3-3’UTR and MUT-DKK3-3’UTR luciferase reporter plasmids (Figure 2C). Luciferase assays demonstrated that the reporter activity was significantly decreased in HCT116 cells transfected with the WT-DKK3-3’UTR construct compared with the parental UTR-less cassette. However, no significant difference in activity was observed in cells transfected with MUT-DKK3-3’UTR, where the predicted miR-92a binding sites had been mutated (Figure 2D).

We next sought to determine whether miR-92a is directly bound to DKK3 mRNA using biotinylated microRNA/mRNA pulldown (Figure 2E). Pulldown of miR-92a itself was confirmed by stem-loop RT-PCR (Supplementary Figure S1C). As anticipated, levels of DKK3 mRNA were markedly elevated in biotin-labeled miR-92a pull-down fraction compared to control, while there was no difference in the levels of HPRT mRNA. (Figure 2F). Consistent with this binding, DKK3 mRNA (Figure 2G) and protein (Figure 2H) levels were significantly decreased compared to those seen in control miR-transfected cells. In order to determine whether Myc regulates DKK3 via miR-92a, we treated Myc overexpressing cells with miR-92a antisense inhibitor. In the presence of this inhibitor, down-regulation of DKK3 by Myc was completely abolished (Figure 2I). As an independent approach, we treated the cells with CHIR99021, which resulted in elevated levels of Myc and reduced levels of DKK3. However, when the experiment was performed in the presence of the miR-92a inhibitor, the suppression of DKK3 by Myc was once again lost (Figure 2J). These findings led us to conclude that miR-92a is the major, if not the sole mediator of DKK3 repression by Myc.

### DKK3 represses Wnt signaling in colon cancer cells and Kras mutated, p53, Apc and Smad4-null organoids

To evaluate the role of DKK3 in the context of WNT activating mutations in CRC, we analyzed the effect of DKK3 overexpression and knockdown on Wnt signaling using HCT116 cells which harbor an activating β-catenin mutation and SK-CO-1 cells which harbor an APC mutation. To induce Wnt signaling, media conditioned by murine Wnt3A-producing or control L cells were used (32). In HCT116 and SK-CO-1 cells upon Wnt3a treatment, LRP6 phosphorylation at Ser 1490, a hallmark of WNT/β-catenin pathway activation, was increased, while the protein level of LRP6 remained unchanged. We also observed higher phosphorylation of GSK3β and increased level of Wnt target genes Axin2 and CCND1. However, in the presence of overexpressed DKK3, all these effects were ablated (Figure 3A, left and right panels). Conversely, knockdown of DKK3 in HCT116 cells increases basal as well as Wnt-induced phospho-LRP6 levels compared to control siRNA-transfected cells (Figure 3A, middle panel). We further confirmed these observations using the β-catenin-responsive TOP FLASH reporter, whose activity was enhanced by Wnt3a, but significantly repressed by DKK3 overexpression in the same cell lines (Figures 3B and D). Conversely, this reporter’s activity was enhanced by DKK3 knockdown (Figure 3C) indicating that DKK3 suppresses Wnt signaling at endogenous levels. To rule out off-target effects of siRNAs, we also used CRISPR-Cas9-mediated genome editing. DKK3-specific short guide RNAs and homology directed repair plasmids containing the puromycin resistance genes were transfected into HCT116 cells and puro-resistant clones were selected. Effective DKK3 knockdown in pooled clones was confirmed by Western Blotting (Supplemental Figure S1E). The effect of DKK3 downregulation on Wnt signaling was analyzed and above, and stronger Wnt-dependent increase in phospo-LRP6 was observed in DKK3 edited cultures (Supplemental Figure S1F). Taken together, these results suggest that DKK3 inhibits Wnt signaling in colorectal cancer cells even in the presence of activating APC and CTNNB1 mutations.

**Figure 3.**
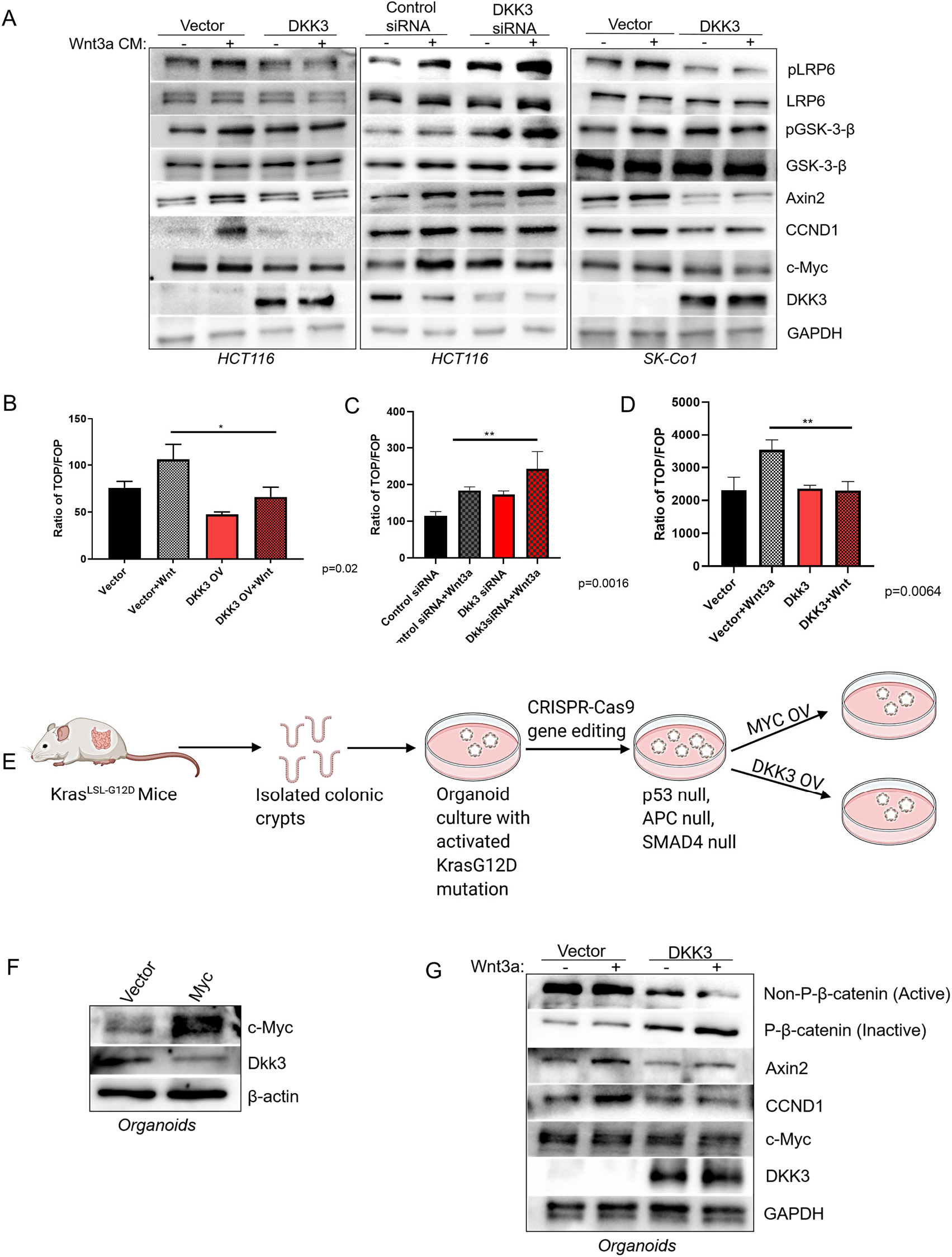
DKK3 represses Wnt signaling in colon cancer cells and tumor organoids. (A) Induction of Wnt signaling by Wnt3a-containing conditioned media. HCT116 and SK-CO-1 cells stably expressing DKK3 or vector alone (left and right panels) and HCT116 cells treated with control and anti-DKK3 siRNA (middle panel) were serum starved and treated with Wnt3a-CM or control L-CM. Total cell lysates was analyzed by immunoblotting using antibodies against pLRP6, pGSK-3β, axin2, CCND1, Myc and DKK3 as indicated. Total LRP6, total pGSK-3β and GAPDH were used as loading control. (B-D) Relative luciferase activities driven by Wnt-responsive TOP FLASH and control FOP FLASH reporters in cells from panel A. (E) Schematic showing the generation of organoids from mouse colonic crypts followed by genome editing. (F) Immunoblotting showing Myc and DKK3 levels in Myc-overexpressing and control organoids. (G) Immunoblotting showing levels of active and inactive β-catenin, axin2, CCND1, Myc and DKK3 in organoids overexpressing vector/DKk3 and treated with 200ng of recombinant Wnt3a for 48 hrs.

For a more genetically defined *ex vivo* model of colon cancer, we used colonic crypts from mice bearing the KrasG12D-LSL mutation (33) to generate transformed organoids. Briefly, using CRISPR/Cas9 genome editing, we further modified them to carry Tp53-, Apc-, and Smad4-null mutations commonly found in human CRC (Figure 3E, Supplemental Figure S2). We additionally engineered them to overexpress retrovirally encoded MYC, which resulted in reduced expression of DKK3 (Figure 3F).

Alternatively, to analyze the effect of DKK3 on Wnt signaling, organoids were stably transduced with control and DKK3-encoding retroviruses and treated with recombinant Wnt3a. We observed that compared to control cultures, DKK3-overexpressing organoids showed higher level of phosphorylated (inactive) β-catenin and reduced levels of non-phosphorylated (active) β-catenin. Consistent with this finding, overexpression of DKK3 blunted induction by Wnt3a of Axin2 and CCND1, its canonical targets (Figure 3G). These results further establish the role of DKK3 as a negative regulator of Wnt signaling in diverse genetic backgrounds.

### Myc enhances Wnt signaling by enabling the miR-92a/DKK3 axis

Since overexpressed Myc down-regulates DKK3, which acts as a Wnt signaling inhibitor, we hypothesized that in addition to being downstream of Wnt in the signal transduction pathway, Myc could reciprocally enhance Wnt signaling and in doing so boost its own expression. To evaluate this scenario, control and Myc-overexpressing HCT116 cells were treated Wnt3a-CM or L-CM. HCT116-Myc cells indeed showed elevated phospho-LRP6 levels (Figure 4A, left panel) and higher levels of Wnt targets Axin2, CCND1 and Myc. Notably, these effects were abolished when HCT116-Myc cells were additionally transfected with either the miR-92 inhibitor or the DKK3 expression cassette (Figure 4A, middle and right panels), fully supporting our hypothesis about the role of the miR-92a/DKK3 axis downstream of Myc. Similar effects were observed using the TOP FLASH reporter assay performed in the same three types of cells (Figure 4B-D). Based on our findings, we proposed a model for Myc and WNT interplay in colorectal cancer cells wherein Myc and Wnt signaling are linked in a feed-forward loop involving suppression of DKK3 expression by miR-92a (Figure 4E).

**Figure 4.**
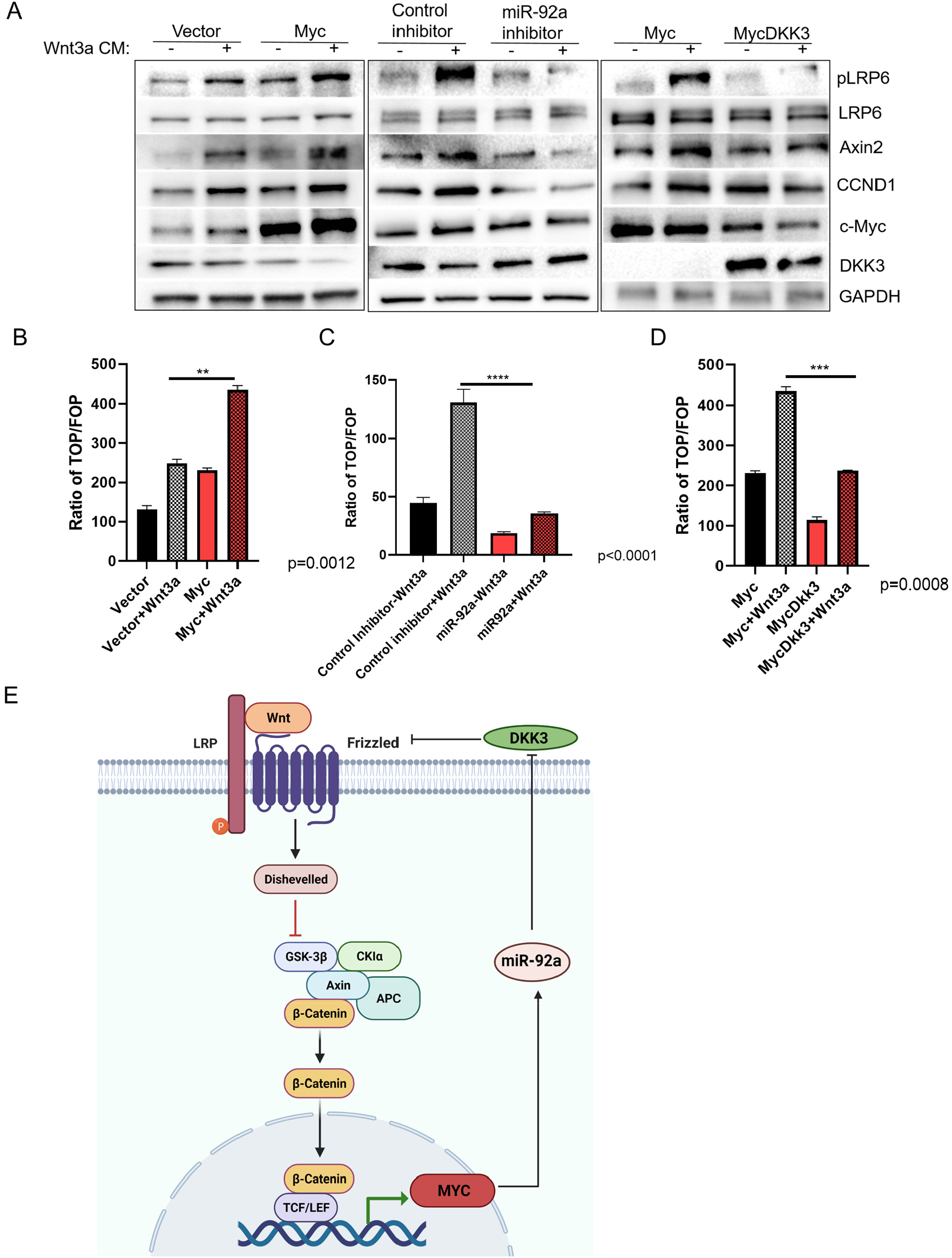
Myc enhances Wnt signaling by enabling the mir-92a/DKK3 axis. (A) Induction of Wnt signaling by Wnt3a-containing conditioned media. HCT116 cells overexpressing vector or Myc (left panel) and HCT116-Myc transfected with either miR-92a inhibitor (middle panel) or DKK3 expression cassette (right panel) were serum starved and treated with Wnt3a-CM or control L-CM. Total cell lysates was analyzed by immunoblotting using antibodies against pLRP6, axin2, CCND1, Myc and DKK3 as indicated. Total LRP6 and GAPDH were used as loading control. (B-D) Relative luciferase activities driven by Wnt-responsive TOP FLASH and control FOP FLASH reporters in cells from panel A (E) The overall model depicting the effect of the Myc->miR-92a-|DKK3 axis on Wnt signaling.

### Myc induces migration and invasion by colon cancer cells via repression of DKK3 and activation of Wnt, resulting in enhanced metastasis

Because Myc is a known regulator of epithelial-mesenchymal transition (EMT) and the motile/invasive phenotype (34), we asked whether DKK3 downregulation plays a role in these processes. We first evaluated the expression of EMT markers in HCT116 cells stably overexpressing Myc. We found that HCT116-Myc cells have increased expression of mesenchymal markers such as N-cadherin and α-smooth muscle actin (α-sma), with a concomitant decrease in the expression of epithelial markers such as E-cadherin. However, restoring DKK3 in these cells noticeably decreased the level of Myc and reversed its effects on EMT markers (Figure 5A).

**Figure 5.**
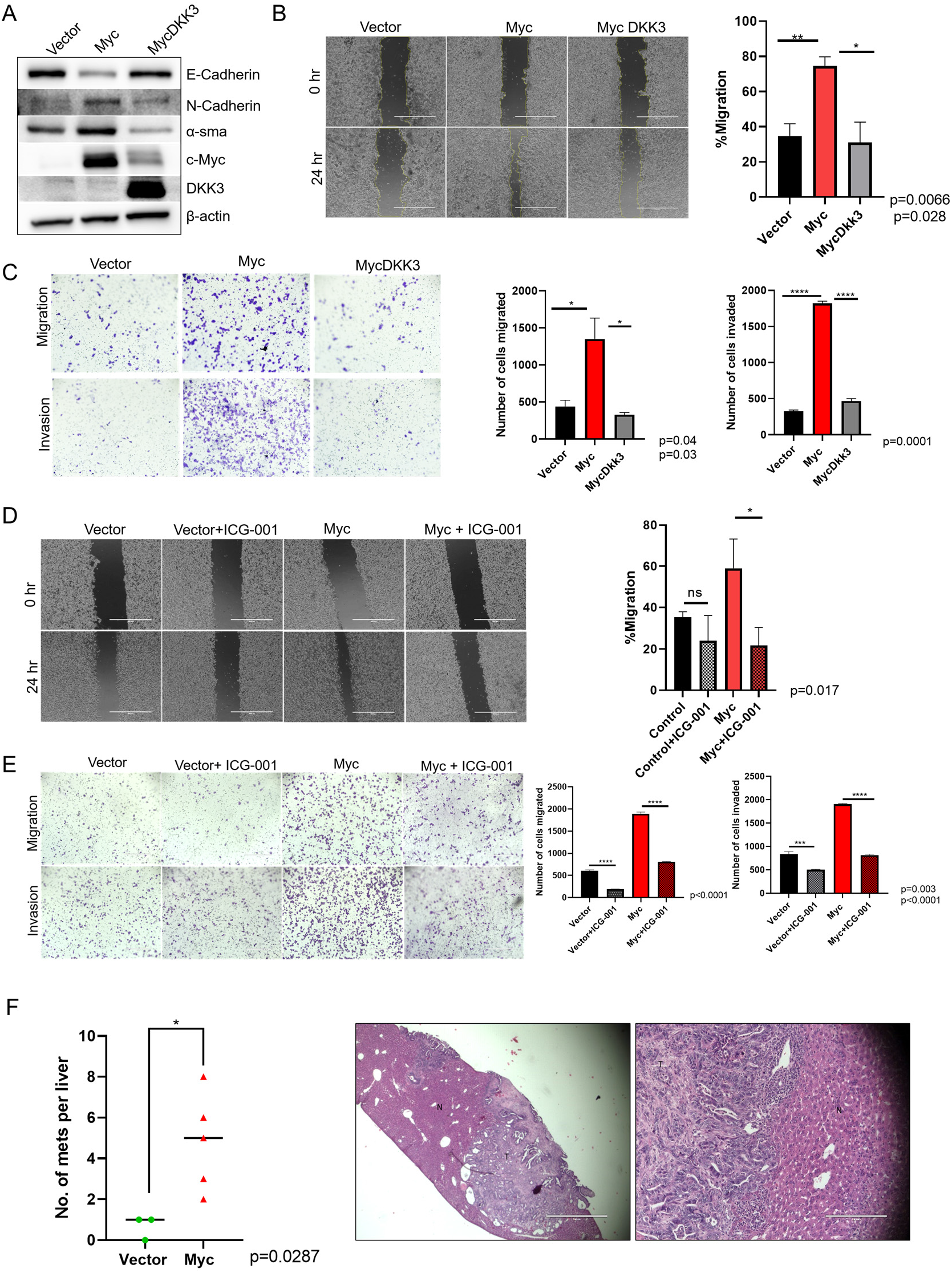
Myc induces migration and invasion by colon cancer cells via repression of DKK3 and activation of Wnt, resulting in enhanced metastasis. (A) Immunoblotting showing expression of EMT markers in HCT116 cells stably expressing vector, Myc, or Myc+DKK3. (B) Migration patterns of the HCT116 cell line derivatives at 24 hrs post wounding. Bars represent the closure of the initial wounded area, in percentages. (C) Migration and invasion potential of the HCT116 cell line derivatives as assessed using the transwell assay, with or without Matrigel, respectively. Numbers of migrated and invaded cells in the lower chambers were plotted, with each bar representing mean ± SEM. All results are representative of three separate experiments each analyzed using Student t-test. (D) Migration patterns of the HCT116 cell line derivatives at 24 hrs post wounding with or without Wnt inhibitor ICG-001. Bars represent the closure of the initial wounded area, in percentages. (E) Migration and invasion potential of the HCT116 cell line derivatives as assessed using the transwell assay, with or without Wnt inhibitor ICG-001. Quantitation of the results was performed as in panel C. (F) Graph showing quantitation of metastatic nodules per liver in mice implanted with vector (n=3) and Myc overexpressing (n=5) organoids on the left. On the right, 4X and 10X micrographs showing H&E staining of Myc-driven mets in liver sections.

Cells undergoing EMT acquire migratory and invasive phenotypes (35). Thus, *in vitro* wound healing (scratch) assays were performed to evaluate the role of Myc in colon cancer cell migration. We determined that HCT116-Myc cells closed the wound twice as efficiently as the control cells, and this effect was fully reversed upon DKK3 re-expression (Figure 5B). Furthermore, Myc overexpression significantly enhanced migration and invasion of HCT116 cells compared with the vector-expressing controls in the regular and matrigel-coated Boyden chambers, respectively. Once again, these effects were reversed in HCT116 Myc+DKK3 cells (Figure 5C). To determine whether Myc-induced migration and invasion are Wnt pathway-dependent, we treated HCT116-Myc cells with ICG-001, a well-validated Wnt inhibitor (36). In both wound healing and Boyden chamber assays, pro-EMT effects of Myc were reversed by ICG-001 (Figure 5D, E). Finally, to establish the effects of Myc on metastasis in vivo, we utilized our syngeneic, orthotopic model of metastatic colorectal carcinoma. The tumoroids with Vector or Myc overexpression were orthotopically transplanted into the cecal walls of syngeneic Bl/6 mice. 6 weeks post-transplant, primary tumors and livers were resected. Immunohistochemical analysis showed that the number of liver metastases was significantly higher in the mice transplanted with Myc overexpressing organoids as compared to vector transplanted mice (Figure 5F). Collectively, these data suggest a role for Myc in the induction of EMT and the associated motile/invasive phenotypes strongly dependent on DKK3 down-regulation and ensuing Wnt pathway activation.

## DISCUSSION

MYC is the central player in CRC, activated either by gene amplification or, by virtue of being a Wnt target gene, via APC deletions or β-catenin activating mutations. However, the full repertoire of Myc-regulated pathways in CRC has not been established. Previous work from other investigators and our own laboratory have demonstrated that many of Myc effects on gene expression are realized through dysregulation of microRNAs [reviewed in (28)], in particular the miR-17-92 cluster microRNA (29,30). Earlier we demonstrated that in CRC Myc-induced miR-17-92 directly targets thrombospondin-1 and other anti-angiogenic factors bringing about the angiogenic switch (20). In addition, miR-17-92 broadly dampens TGF-β signaling (e.g., via targeting of SMAD4) (21,37,38), which is often a barrier to CRC neovascularization (22).

Here we demonstrate that another miR-17-92 direct target is DKK3. While this cluster encodes six distinct microRNAs (miR-17, -18a, -19a, -20a, -19b, and -92a), miR-92a by far is the most abundant (39), serves as a proposed biomarker for early detection of CRC (40,41) and according to some reports is associated with worse prognosis in CRC patients (42). Notably, ectopic overexpression miR92a has been shown to promote stem cell characteristics, proliferation and migration of CRC cells by down-regulating multiple components of the Wnt pathway including, but not limited to, DKK3 (43,44). However, neither the oncogenic mechanism of miR-92a overexpression nor its direct binding of DKK3 mRNA have been established. Here we demonstrate direct binding of miR-92a to the DKK3 transcript, establishing DKK3 as a *bona fide* miR-92a target. We also show that miR-92 mediates repression of DKK3 by Myc via the seed homology sequence in DKK3 3’UTR.

The mir-92a-DKK3 connection was recently reported to contribute to osteosarcoma cell proliferation by a yet to be determined mechanism (45). Similarly, the functional relationship between DKK3 and another member of the Myc family N-Myc was previously reported to exist in B-ALL and neuroblastoma, but either the underlying microRNA-based mechanism (46) or the connection to Wnt signaling (47) were left unexplored. In contrast, our work places the mir-92a-DKK3 axis in the middle of the feedforward loop by which Myc levels and Wnt signaling sustain each other (Figure 4E). Surprisingly, this interplay unfolds in cells where Wnt signaling is thought to be constitutively active because of acquired mutations in this pathway. Furthermore, as a consequence of hyperactive Wnt signaling, Myc is able to induce expression of EMT markers and promote migration and invasion of colon cancer cells by a DKK3 downregulation-dependent mechanism. This effect of Myc was also observed in our syngeneic, orthotopic model of metastatic colorectal carcinoma where mice implanted with Myc overexpressing tumoroids showed enhanced liver metastasis.

The role of DKK3 in Wnt signaling has been controversial and highly context dependent. For example, unlike DKK1, DKK3 is unable to inhibit Wnt signaling in Xenopus embryos to set up secondary axis induction (48). More recent work demonstrated that DKK3 promotes Wnt signaling in Muller glia MIOM1 and HEK293 cell lines (49), inhibits it in pheochromocytoma PC12 and osteocarcinoma Saos-2 cells (50,51), and has no effect in LNCaP prostate cancer cells (52). From broader studies with multiple cell lines, the consensus is emerging that DKK3 generally antagonizes Wnt signaling and cell proliferation in lung (53), breast (54), and cervical (55) cancers. The main caveat is that these studies were performed in cells presumably lacking Wnt pathway mutations. We found that manipulating DKK3 levels profoundly affects Wnt signaling in murine and human colon cancer cells with pre-existing CTNNB1 and APC mutations. This observation is conceptually similar to the finding that restoration of secreted frizzled-related proteins (SFRP) expression in colon cancer cells attenuates Wnt signaling despite the presence of APC/CTNNB1 mutations (56). Interestingly, SFRP1 and DKK1 are repressed by Myc in mammary epithelial cell lines (57) suggesting that inhibition of secreted Wnt antagonist might be a common theme in Myc-driven oncogenic programs. This could be a potent complement to the reported ability to Myc to activate transcription (58) and potentiate translation (59) of several key Wnt signaling components including Lef1. More broadly speaking, Myc effects on Wnt signaling highlight the importance of “secondary” hyperactivating events and help explain the surprising pattern of seemingly redundant but still co-occurring mutations in human CRC.

## ACKNOWLEDGEMENTS

The authors thank members of their laboratories for many helpful discussions. Initial experiments on DKK3 downregulation in the ATT group were performed by Grace Tan. Wnt3a-overexpressing mouse L-cells were a kind gift from Patrick Viatour (CHOP). This work was supported by NIH grant R01 CA196299 and funds from the Colorectal Cancer Translational Center of Excellence of the Abramson Cancer Center at the University of Pennsylvania. All the schematics in the manuscript have been created with BioRender.com

---

**Supplemental Figure 1.**
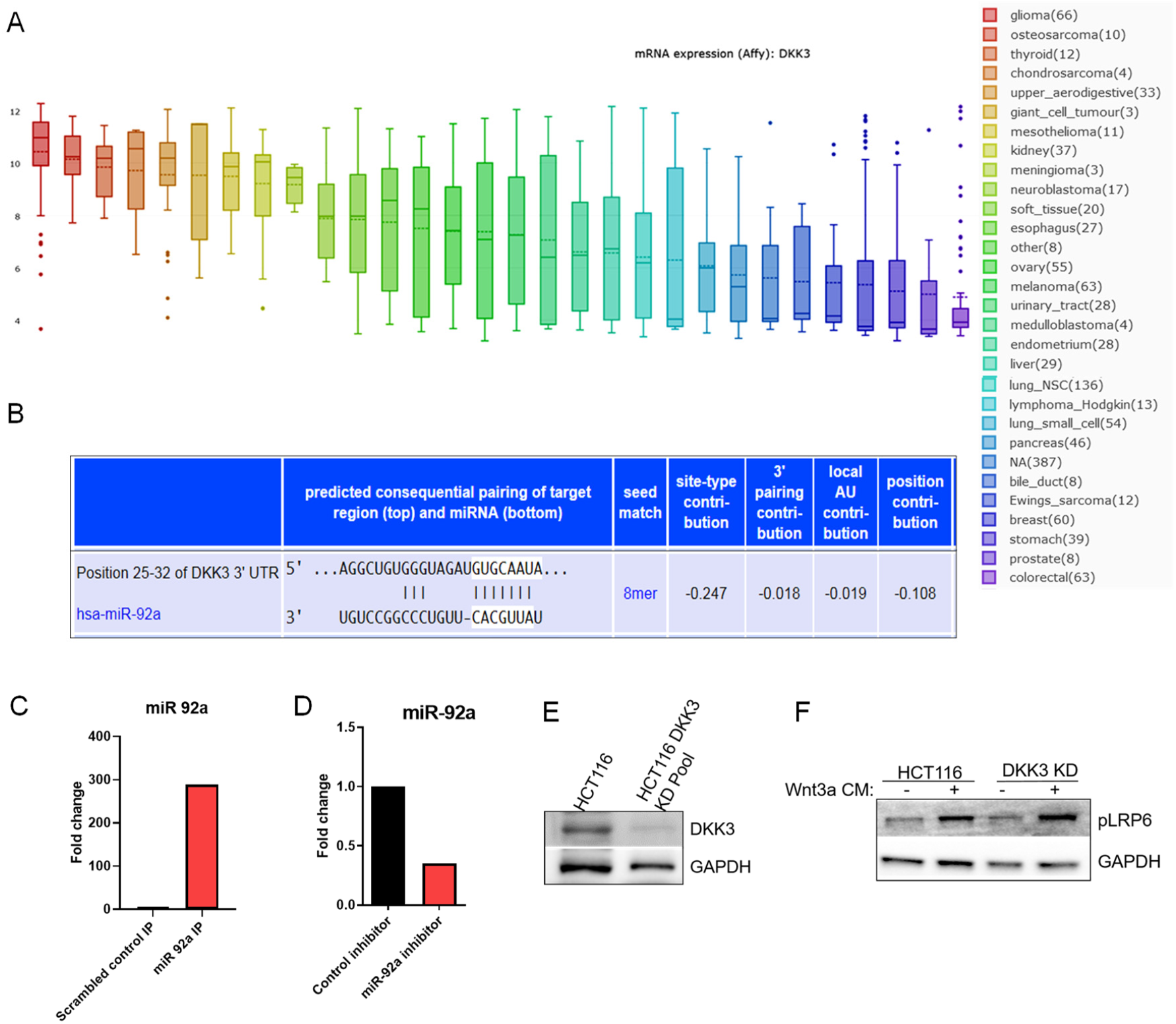
DKK3 and Wnt signaling in CRC cells. (A) DKK3 mRNA expression across various solid cancer types profiled in the Broad Cell Line Encyclopedia. B) Predicted targeting of DKK3 3’UTR by miR-92a. C-D) Bar graph represents Log2 transformed ratio of miR92a expression normalized to RNU6B as determined by miRNA qRT-PCR analysis. E) Immunoblotting showing levels of DKK3 in CRISPR/Cas9 edited DKK3 knockdown pooled clones. GAPDH is used as loading control. F) HCT116 parental cells and DKK3 CRISPR/Cas9 Knockdown pooled clones were treated with Wnt3a-CM or control L-CM. Total cell lysates was analyzed by immunoblotting using antibodies against pLRP6 and GAPDH as indicated.

**Supplemental Figure 2.**
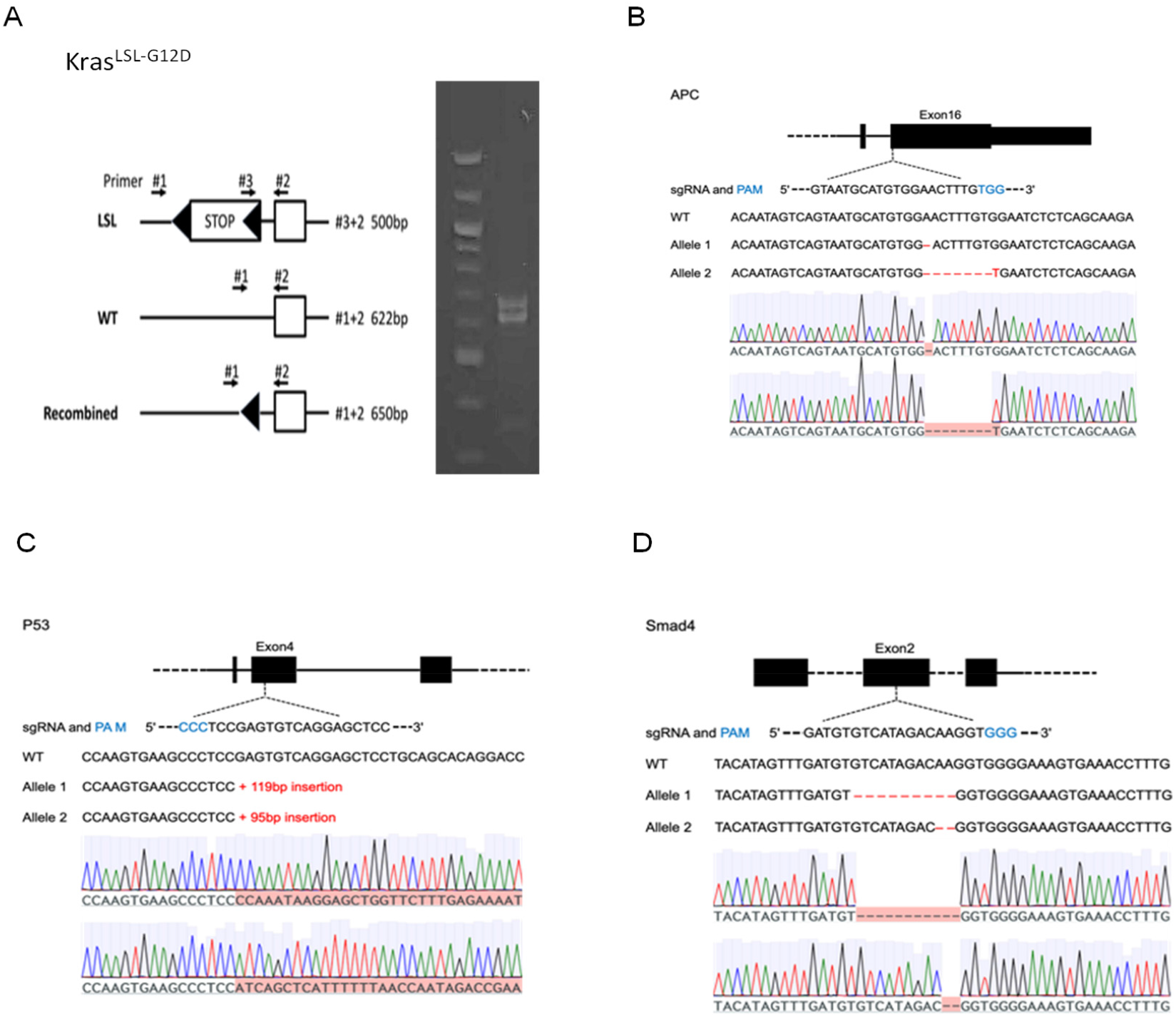
Derivation of murine colorectal tumoroids. (A-D) colonic organoids from a KrasLSL-G12D mouse were transient transfected with Salk-Cre and PGK-puro plasmids, after being selected with puromycin, the organoids were further transient transfected with plasmids that express sgRNAs targeting APC, SMAD4, or TP53, and selected by removing R-spondin, Noggin and adding Nutlin-3 in the culture media. Subcloned organoids with mutations confirmed by TA clone and Sanger sequencing were picked and used in the downstream assays. (A) Kras G12D conditional PCR showed recombined LSL allele at 650bp and WT allele at ∼622bp. (B-D) Strategy to generate indicated mutations and targeted loci, and sequences of the mutated gene alleles.

## REFERENCES

1. Siegel RL, Miller KD, Jemal A. Cancer statistics, 2020. CA Cancer J Clin 2020;70:7–30

2. Sansom OJ, Meniel VS, Muncan V, Phesse TJ, Wilkins JA, Reed KR, et al. Myc deletion rescues Apc deficiency in the small intestine. Nature 2007;446:676–9

3. He TC, Sparks AB, Rago C, Hermeking H, Zawel L, da Costa LT, et al. Identification of c-MYC as a target of the APC pathway. Science 1998;281:1509–12

4. MacDonald BT, Tamai K, He X. Wnt/beta-catenin signaling: components, mechanisms, and diseases. Dev Cell 2009;17:9–26

5. Stamos JL, Weis WI. The beta-catenin destruction complex. Cold Spring Harb Perspect Biol 2013;5:a007898

6. Clevers H, Nusse R. Wnt/beta-catenin signaling and disease. Cell 2012;149:1192–205

7. Bienz M, Clevers H. Linking colorectal cancer to Wnt signaling. Cell 2000;103:311–20

8. van de Wetering M, Sancho E, Verweij C, de Lau W, Oving I, Hurlstone A, et al. The beta-catenin/TCF-4 complex imposes a crypt progenitor phenotype on colorectal cancer cells. Cell 2002;111:241–50

9. Morin PJ, Sparks AB, Korinek V, Barker N, Clevers H, Vogelstein B, et al. Activation of β-Catenin-Tcf Signaling in Colon Cancer by Mutations in β-Catenin or APC. Science 1997;275:1787–90

10. Sparks AB, Morin PJ, Vogelstein B, Kinzler KW. Mutational analysis of the APC/beta-catenin/Tcf pathway in colorectal cancer. Cancer Res 1998;58:1130–4

11. Mao B, Wu W, Li Y, Hoppe D, Stannek P, Glinka A, et al. LDL-receptor-related protein 6 is a receptor for Dickkopf proteins. Nature 2001;411:321–5

12. Niehrs C. Function and biological roles of the Dickkopf family of Wnt modulators. Oncogene 2006;25:7469–81

13. Nakamura RE, Hackam AS. Analysis of Dickkopf3 interactions with Wnt signaling receptors. Growth factors (Chur, Switzerland) 2010;28:232–42

14. Mao B, Wu W, Davidson G, Marhold J, Li M, Mechler BM, et al. Kremen proteins are Dickkopf receptors that regulate Wnt/beta-catenin signalling. Nature 2002;417:664–7

15. Andersson-Rolf A, Fink J, Mustata RC, Koo BK. A video protocol of retroviral infection in primary intestinal organoid culture. Journal of visualized experiments : JoVE 2014:e51765

16. Lanauze CB, Sehgal P, Hayer K, Torres-Diz M, Pippin JA, Grant SFA, et al. Colorectal cancer-associated Smad4 R361 hotspot mutations boost Wnt/β-catenin signaling through enhanced Smad4-LEF1 binding. Mol Cancer Res 2021:molcanres.0721.2020

17. Orom UA, Lund AH. Isolation of microRNA targets using biotinylated synthetic microRNAs. Methods (San Diego, Calif) 2007;43:162–5

18. Muzny DM, Bainbridge MN, Chang K, Dinh HH, Drummond JA, Fowler G, et al. Comprehensive molecular characterization of human colon and rectal cancer. Nature 2012;487:330–7

19. Hoadley KA, Yau C, Hinoue T, Wolf DM, Lazar AJ, Drill E, et al. Cell-of-Origin Patterns Dominate the Molecular Classification of 10,000 Tumors from 33 Types of Cancer. Cell 2018;173:291–304.e6

20. Dews M, Homayouni A, Yu D, Murphy D, Sevignani C, Wentzel E, et al. Augmentation of tumor angiogenesis by a Myc-activated microRNA cluster. Nature genetics 2006;38:1060–5

21. Dews M, Fox JL, Hultine S, Sundaram P, Wang W, Liu YY, et al. The Myc-mir-17∼92 axis blunts TGFβ signaling and production of multiple TGFβ-dependent antiangiogenic factors. Cancer Res 2010;70:8233–46

22. Dews M, Tan GS, Hultine S, Raman P, Choi J, Duperret EK, et al. Masking epistasis between MYC and TGF-beta pathways in antiangiogenesis-mediated colon cancer suppression. J Natl Cancer Inst 2014;106:dju043

23. Chaidos A, Caputo V, Gouvedenou K, Liu B, Marigo I, Chaudhry MS, et al. Potent antimyeloma activity of the novel bromodomain inhibitors I-BET151 and I-BET762. Blood 2014;123:697–705

24. Harrington CT, Sotillo E, Robert A, Hayer KE, Bogusz AM, Psathas J, et al. Transient stabilization, rather than inhibition, of MYC amplifies extrinsic apoptosis and therapeutic responses in refractory B-cell lymphoma. Leukemia 2019;33:2429–41

25. Farrell AS, Sears RC. MYC Degradation. Cold Spring Harbor Perspectives in Medicine 2014;4:a014365

26. Dejure FR, Royla N, Herold S, Kalb J, Walz S, Ade CP, et al. The MYC mRNA 3’-UTR couples RNA polymerase II function to glutamine and ribonucleotide levels. The EMBO journal 2017;36:1854–68

27. Zhang J, Chen S, Zhang W, Zhang J, Liu X, Shi H, et al. Human differentiation-related gene NDRG1 is a Myc downstream-regulated gene that is repressed by Myc on the core promoter region. Gene 2008;417:5–12

28. Psathas JN, Thomas-Tikhonenko A. MYC and the art of microRNA maintenance. Cold Spring Harb Perspect Med 2014;4:a014175

29. O’Donnell KA, Wentzel EA, Zeller KI, Dang CV, Mendell JT. c-Myc-regulated microRNAs modulate E2F1 expression. Nature 2005;435:839–43

30. He L, Thomson JM, Hemann MT, Hernando-Monge E, Mu D, Goodson S, et al. A microRNA polycistron as a potential human oncogene. Nature 2005;435:828–33

31. Agarwal V, Bell GW, Nam J-W, Bartel DP. Predicting effective microRNA target sites in mammalian mRNAs. eLife 2015;4:e05005

32. Willert K, Brown JD, Danenberg E, Duncan AW, Weissman IL, Reya T, et al. Wnt proteins are lipid-modified and can act as stem cell growth factors. Nature 2003;423:448–52

33. Jackson EL, Willis N, Mercer K, Bronson RT, Crowley D, Montoya R, et al. Analysis of lung tumor initiation and progression using conditional expression of oncogenic K-ras. Genes & development 2001;15:3243–8

34. Wolfer A, Ramaswamy S. MYC and metastasis. Cancer Res 2011;71:2034–7

35. Tsai JH, Yang J. Epithelial-mesenchymal plasticity in carcinoma metastasis. Genes & development 2013;27:2192–206

36. McMillan M, Kahn M. Investigating Wnt signaling: a chemogenomic safari. Drug discovery today 2005;10:1467–74

37. Fox JL, Dews M, Minn AJ, Thomas-Tikhonenko A. Targeting of TGFβ signature and its essential component CTGF correlates with improved survival in glioblastoma. RNA 2013;19

38. Mestdagh P, Bostrom AK, Impens F, Fredlund E, Van PG, De AP, et al. The miR-17-92 microRNA cluster regulates multiple components of the TGF-β pathway in neuroblastoma. Mol Cell 2010;40:762–73

39. Tsuchida A, Ohno S, Wu W, Borjigin N, Fujita K, Aoki T, et al. miR-92 is a key oncogenic component of the miR-17–92 cluster in colon cancer. Cancer Science 2011;102:2264–71

40. Schee K, Boye K, Abrahamsen TW, Fodstad Ø, Flatmark K. Clinical relevance of microRNA miR-21, miR-31, miR-92a, miR-101, miR-106a and miR-145 in colorectal cancer. BMC Cancer 2012;12:505

41. Wu CW, Ng SSM, Dong YJ, Ng SC, Leung WW, Lee CW, et al. Detection of miR-92a and miR-21 in stool samples as potential screening biomarkers for colorectal cancer and polyps. Gut 2012;61:739–45

42. Zhou T, Zhang G, Liu Z, Xia S, Tian H. Overexpression of miR-92a correlates with tumor metastasis and poor prognosis in patients with colorectal cancer. International journal of colorectal disease 2013;28:19–24

43. Zhang GJ, Li LF, Yang GD, Xia SS, Wang R, Leng ZW, et al. MiR-92a promotes stem cell-like properties by activating Wnt/β-catenin signaling in colorectal cancer. Oncotarget 2017;8:101760–70

44. Chen E, Li Q, Wang H, Yang F, Min L, Yang J. MiR-92a promotes tumorigenesis of colorectal cancer, a transcriptomic and functional based study. Biomedicine & pharmacotherapy = Biomedecine & pharmacotherapie 2018;106:1370–7

45. Yu H, Song H, Liu L, Hu S, Liao Y, Li G, et al. MiR-92a modulates proliferation, apoptosis, migration, and invasion of osteosarcoma cell lines by targeting Dickkopf-related protein 3. Bioscience reports 2019;39

46. Kong D, Zhao L, Sun L, Fan S, Li H, Zhao Y, et al. MYCN is a novel oncogenic target in adult B-ALL that activates the Wnt/β-catenin pathway by suppressing DKK3. Journal of cellular and molecular medicine 2018;22:3627–37

47. De Brouwer S, Mestdagh P, Lambertz I, Pattyn F, De PA, Westermann F, et al. Dickkopf-3 is regulated by the MYCN-induced miR-17-92 cluster in neuroblastoma. International journal of cancer 2012;130:2591–8

48. Krupnik VE, Sharp JD, Jiang C, Robison K, Chickering TW, Amaravadi L, et al. Functional and structural diversity of the human Dickkopf gene family. Gene 1999;238:301–13

49. Nakamura RE, Hunter DD, Yi H, Brunken WJ, Hackam AS. Identification of two novel activities of the Wnt signaling regulator Dickkopf 3 and characterization of its expression in the mouse retina. BMC cell biology 2007;8:52

50. Caricasole A, Ferraro T, Iacovelli L, Barletta E, Caruso A, Melchiorri D, et al. Functional characterization of WNT7A signaling in PC12 cells: interaction with A FZD5 x LRP6 receptor complex and modulation by Dickkopf proteins. The Journal of biological chemistry 2003;278:37024–31

51. Hoang BH, Kubo T, Healey JH, Yang R, Nathan SS, Kolb EA, et al. Dickkopf 3 inhibits invasion and motility of Saos-2 osteosarcoma cells by modulating the Wnt-beta-catenin pathway. Cancer Res 2004;64:2734–9

52. Kawano Y, Kitaoka M, Hamada Y, Walker MM, Waxman J, Kypta RM. Regulation of prostate cell growth and morphogenesis by Dickkopf-3. Oncogene 2006;25:6528–37

53. Yue W, Sun Q, Dacic S, Landreneau RJ, Siegfried JM, Yu J, et al. Downregulation of Dkk3 activates β-catenin/TCF-4 signaling in lung cancer. Carcinogenesis 2008;29:84–92

54. Wang XY, Yin Y, Yuan H, Sakamaki T, Okano H, Glazer RI. Musashi1 modulates mammary progenitor cell expansion through proliferin-mediated activation of the Wnt and Notch pathways. Mol Cell Biol 2008;28:3589–99

55. Lee EJ, Jo M, Rho SB, Park K, Yoo YN, Park J, et al. Dkk3, downregulated in cervical cancer, functions as a negative regulator of beta-catenin. IntJ Cancer 2009;124:287–97

56. Suzuki H, Watkins DN, Jair KW, Schuebel KE, Markowitz SD, Chen WD, et al. Epigenetic inactivation of SFRP genes allows constitutive WNT signaling in colorectal cancer. Nature genetics 2004;36:417–22

57. Cowling VH, D’Cruz CM, Chodosh LA, Cole MD. c-Myc transforms human mammary epithelial cells through repression of the Wnt inhibitors DKK1 and SFRP1. Mol Cell Biol 2007;27:5135–46

58. Hao YH, Lafita-Navarro MC, Zacharias L, Borenstein-Auerbach N, Kim M, Barnes S, et al. Induction of LEF1 by MYC activates the WNT pathway and maintains cell proliferation. Cell communication and signaling : CCS 2019;17:129

59. Posternak V, Ung MH, Cheng C, Cole MD. MYC Mediates mRNA Cap Methylation of Canonical Wnt/β-Catenin Signaling Transcripts By Recruiting CDK7 and RNA Methyltransferase. Mol Cancer Res 2017;15:213–24

